# Microgravity stimulates network activity of 3D neuronal spheroids in an acoustic trap

**DOI:** 10.1101/2024.07.03.601873

**Authors:** Lecoq Pierre-Ewen, Viraye Guillaume, Dupuis Chloé, Benoit-Gonin Xavier, Aider Jean-Luc, Peyrin Jean-Michel

## Abstract

Among biological models, cell culture constitutes an important paradigm that allows rapid examination of cell phenotype and behavior. While cell cultures are classically grown on a 2D substrate, the recent development of organoid technologies represents a paradigmatic shift in biological experimentation as they pave the way for the reconstruction of minimalist organs in 3D. Manipulating these 3D cell assemblies represents a considerable challenge. While there is growing interest in studying the behavior of cells and organs in the space environment, manipulating 3D cultures in microgravity remains a challenge. But with cellular research underway aboard the International Space Station (ISS), optimizing techniques for handling 3D cellular assemblies is essential. Here, in order to cultivate 3D models of spheroids in microgravity, we developed and used an acoustic bioreactor to trap levitating cellular organoids in a liquid cell culture medium. Indeed, in a Bulk Acoustic Wave (BAW) resonator, spherical objects, such as cells, can be maintained in an equilibrium position, inside a resonant cavity, away from the walls. In the acoustic levitation plane, gravity is counterbalanced by the acoustic radiation force (ARF) making it possible to maintain an object even in weightlessness. A dedicated setup was designed and built to perform live calcium imaging during parabolic flights. During a parabolic flight campaign, we were able to monitor the calcium activity of 3D neural networks trapped in an acoustic field during changes in gravity during different parabolas. Our results clearly indicate a change in calcium activity associated with variations in gravity.

## Introduction

Microgravity impairs physiology and health of the astronauts. When living on the International Space Station (ISS), many physiological changes are observed, such as bone density loss, muscle atrophy, cardiovascular and hemodynamic alterations, metabolic changes, endocrine disorders, and sleep irregularities. It also causes vestibular and ocular structural dysfunction induced by increased intracranial and intraocular pressure, leading to SANS (Spaceflight-associated neuroocular syndrome) (1, 2). While, these result from prolonged exposure to microgravity and body fluids reshuffle, most of the syndromes are reversible upon return on earth. Yet, question are raised on the consequences of longer stay and/or exposure to space radiations that may cause more prolonged deficits.

From a neurological perspective, medical studies of astronauts have also shown that spaceflight has short-term (motion sickness, ventricular enlargement and local intracranial pressure, vestibular dysfunction and SANS) that raises fears about long-term effects on the brain (increased risk of brain tumors, premature aging, accelerated neurodegeneration) (3, 4). Extensive fMRI analysis of astronaut brain structure have recently suggested that astronauts experience subtle but significant changes in the brain connectome (5). These phenomena are either related to microgravity itself, via mechanical unloading, or to intracranial fluid redistribution, or to space radiation.

With the growing interest in the space industry and preparation for long-term stays in the space environment, it is of paramount importance to understand the mechanisms underlying space-specific physiological adaptations and to determine whether or not the space environment triggers pathophysiological conditions.

While physiological studies provide valuable insight into the broader effects of microgravity on organ systems, cellular studies can help elucidate the underlying mechanisms driving these changes. However, conducting cellular studies in microgravity poses significant challenges. Indeed, carrying out these experiments in space require extensive planning, specialized equipment and logistical coordination. Ground-based methods range from experiments simulating microgravity in the laboratory to specific environments creating transient phases of microgravity, such as in rockets, drop towers or parabolic flights. They offer shorter experiment durations, ranging from a few seconds (drop towers, parabolic flights) to a few minutes (rockets) (6). Experiments on the International Space Station (ISS) or in automated space laboratories such as Cube Sats offer the possibility of longer-term studies, but require the development of dedicated tools allowing the manipulation of living cells in a zero-G environment (7, 8).

Although cell biology experiments are regularly carried out in a constrained environment such as the ISS, they are limited due to technical difficulties. Most cellular studies rely on the use of a flat 2D environment, which allows observation by microscopy. However, the evaluation of biological processes in more complex models such as 3D organoïd cultures is limited to analysis on Earth, before and after the stay in space. Manipulating objects, including biological models, in microgravity presents unique challenges and opportunities due to the lack of sedimentation that allows cell culture on Earth. Indeed, in the absence of gravity, objects float in the environment and can drift at the slightest contact, thus complicating precise positioning. Additionally, it is difficult to maintain the orientation and position of tools and objects without a stable reference point (6, 9).

To solve these problems, several methods have been proposed, from trapping cells in bioactive gels (10) to magnetic labeling of living cells allowing levitation in a paramagnetic liquid cell culture medium (11). Acoustic levitation is another technology allowing the manipulation and trapping of microscopic objects such as cells or particles, without mechanical contact. It has already been used during parabolic flights to study the self-propulsion of acoustically levitating metal nanorods in microgravity (12). Acoustic levitation relies on the creation of a stationary acoustic wave inside a resonant cavity, also called Bulk Acoustic Wave (BAW). Variations in acoustic pressure lead to the creation of an acoustic radiation force (ARF). The axial component of the ARF pushes all suspended objects toward the acoustic pressure nodes (minimum pressure regions) of the standing wave. Objects can then be trapped in pressure nodes, also called acoustic levitation planes, where they can be held for as long as necessary in a stable position. The magnitude of this force depends on the acoustic wave properties (frequency, amplitude) as well as of the properties of the objects and the surrounding environment (acoustic contrast). Objects are also trapped radially by the transverse component of the ARF, which is approximately ten times weaker than the axial component (13). It has been demonstrated that BAW resonators can be used to structure, levitate and culture mammalian cells and enable the self-organization of large cellular spheroids (14, 15), in a dynamic manner (16, 17) making acoustic levitation an interesting alternative for manipulating and growing cells in microgravity.

In this study, an original experimental setup was developed. It includes a fluorescence microscope and an acoustofluidic bioreactor allowing the levitation and trapping of large cellular spheroids composed of active neurons. This new experimental platform has been validated by recording the neuronal activity of acoustically trapped neuronal spheroids during a parabolic flight (177th parabolic flight campaign funded by the CNES). It demonstrates that acoustic levitation is a powerful tool for trapping active living objects in a microgravity environment, thereby enabling the microscopic study of microgravity-induced variation in neuronal activity. The experimental measurements indicate that the microgravity environment modifies the calcium activity of neuronal networks. They also show that acoustic levitation could prove to be a particularly interesting tool for all biological experiments involving 3D cell cultures such as cellular spheroids or organoids.

## Results

### Acoustic trapping in micro or hyper-gravity

Performing microscopy imaging during parabolic flights presents unique challenges due to abrupt gravity changes and high noise and vibration generated by aircraft engines. These conditions can interfere with such sensitive imaging equipment and make the data collection difficult. In order to consider live imaging of active cellular objects such as neuronal spheroids in such a harsh environment, efficient and robust trapping of the living cellular objects to be imaged is required. In microgravity, while traditional methods of holding and manipulating delicate structures may fail, acoustic levitation could prove an easy way to hold living objects in a stable position in a liquid culture medium, allowing precise observations and interventions without damaging the objects by mechanical contacts. Indeed, the acoustic radiation force should be able to counterbalance the weight of the object and keep it in acoustic levitation in both micro- and hyper-gravity.

The acoustofluidic chip consists of a PDMS (polydimethylsiloxane) body bonded to a microscope cover glass. The PDMS material was chosen because it is biocompatible and gas permeable, making it ideal for cell culture (18). It is also transparent, allowing optical access, and is easy to process. A 2 *MHz* ultrasonic transducer is in direct contact with the PDMS body, closing the cylindrical cavity (Fig. 1). The microscope cover glass, facing the transducer, acts as an efficient acoustic reflector and allows optical access to visualize objects inside the resonant cavity using an inverted microscope.

**Fig. 1.**
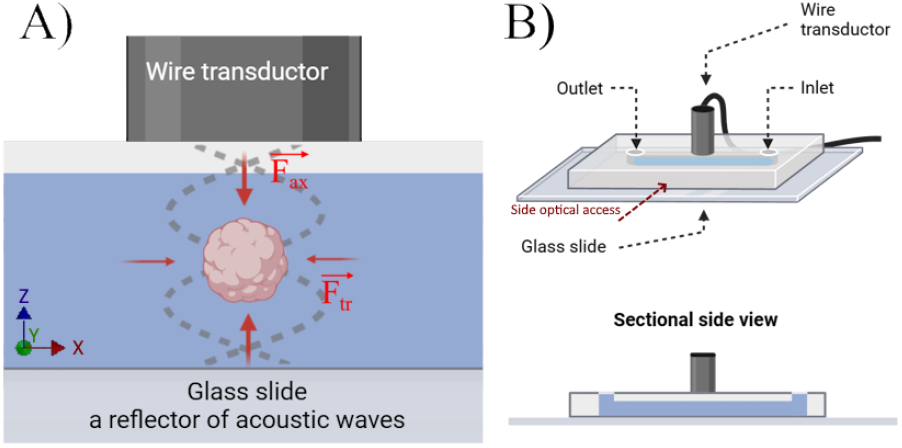
Sketch of the acoustic chip. (A) Side view of the acoustofluidic chip showing the standing acoustic wave inside the PDMS cavity. The typical frequency is 2 *MHz*, corresponding to an acoustic wavelength *λ*_*ac*_ *≈* 375 *µm*. The axial force 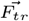 is 10 times stronger than the transverse force 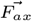.The acoustic wave propagates in water and the reflector is a 1 *mm* thick glass slide. (B) Top view and side view of the PDMS chip with the acoustic transducer on top.

Two acoustofluidics chips and two different setups were used. The first is a 5 *mm* high multinode chip, which allows simple observation from the side, allowing to easily track the objects trapped under acoustic levitation. It will be used to validate the acoustic trapping of large objects even during the changes of gravity. The second is a 700 *µm* high single-node chip. It allows an easy observation from under the chip and is well adapted to fluorescence microscopy with an inverted microscope.

The robustness of acoustic trapping was first tested using large beads of diameter *d*_*p*_ = 300 *µm*, close to the dimension of a cell spheroid. The multi-node acoustofluidic chip was used with an acoustic frequency of 2 *MHz* and a voltage amplitude of 5 *V*. To describe the displacement of large scale objects during a parabola the displacements of beads in acoustic levitation, in ten different levitation planes, were recorded in the *X/Z* plane, in the reference frame of the plane. A typical result is shown in Fig. 2. The Fig. 2A shows the full trajectory of a bead during gravity changes and the evolution of the corresponding *X* and *Z* position as a function of time. The first observation is that the displacements along the *Z* direction are ten times smaller than along the *X* direction. This result is confirmed when looking at the averaged maximal displacements for ten different particles shown in Fig. 2B. The maximum displacement along the *X* axis is at most a third of the diameter of the bead while it is less than 3% of the bead diameter for the *Z* displacement. This result can be explained by the fact that the axial component of the ARF is indeed one order of magnitude larger than its transverse component. It confirms that the acoustic trapping is indeed strong enough and can maintain large objects in acoustic levitation even during the hyper-gravity phase.

**Fig. 2.**
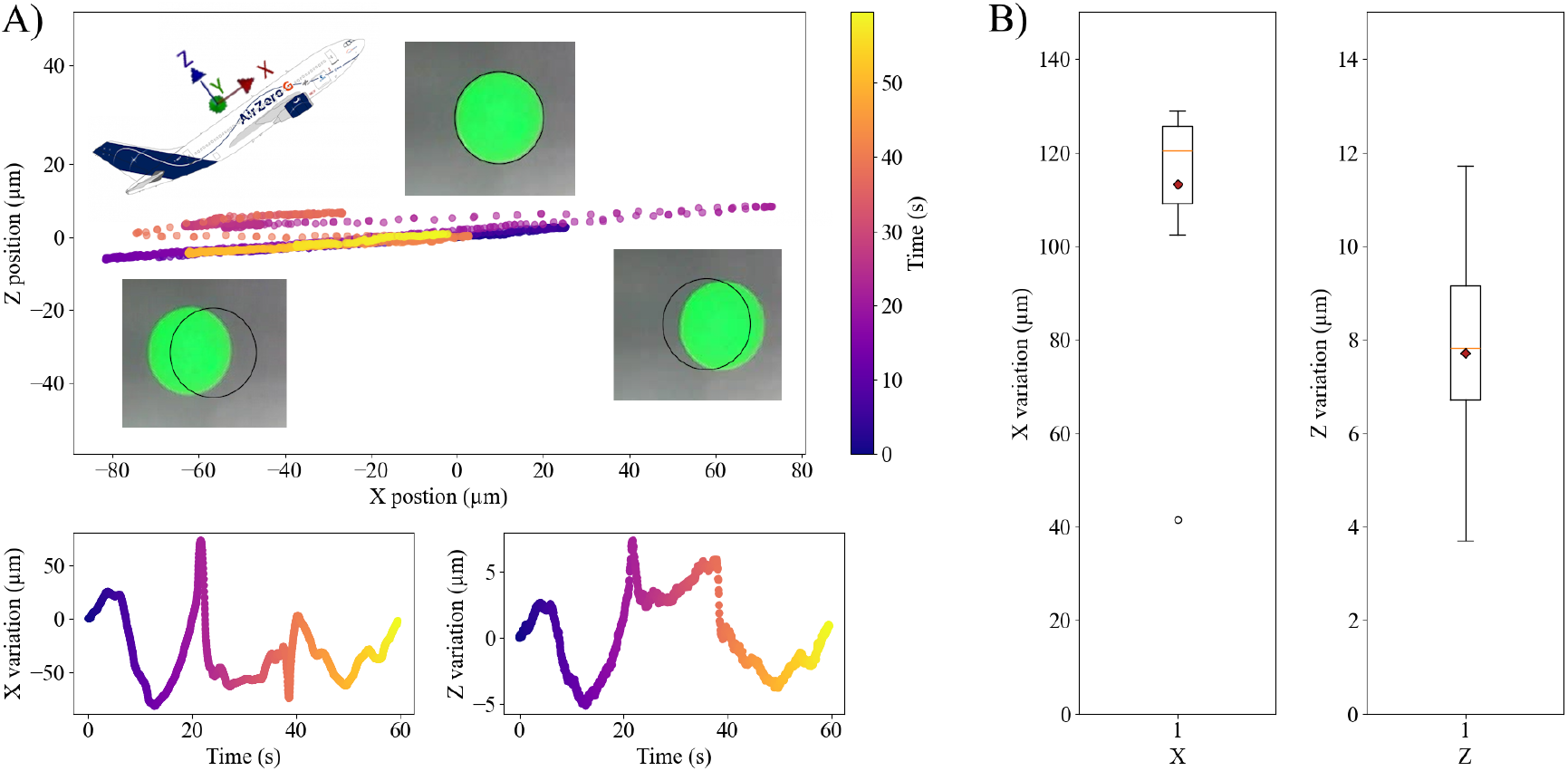
Description of acoustic trapping. (A) Typical trajectory, in the *X/Z* plane, of a 300*µm* bead trapped in acoustic levitation during a parabola. The two secondary graphs show the corresponding time evolution of the position of the bead along these axes. Positions were recorded using a DinoLite digital camera. Images were then processed using a Python code to track the position using the OpenCv library (19). The color gradient is indicative of the progression of time. Each picture is a view of the bead at an extreme position along the *X* axis. (B) Maximum displacement on both *X* (107 *±* 8*µm*) and *Z* axis (7.72 *±* 2.62*µm*) for ten 300*µm* beads trapped in acoustic levitation during a parabola. Red square is the mean value while orange bar is the median. The lines correspond to the first and third quartiles.

These results confirm that the acoustofluidic chip can indeed be used to levitate and trap even large cell spheroids and should subsequently enable live recording of fluorescence microscopy of living cell spheroids.

### Monitoring calcium activity of spheroids

Calcium imaging is a valuable technique for studying the influence of gravity variations on a neural network during parabolic flights. This method is particularly useful because it can capture rapid changes in neuronal activity, typically in ten seconds or less, making it ideal for the brief gravity phases of these flights. Calcium imaging tracks fluorescence fluctuations of many neurons simultaneously, with typical signal frequencies around a few Hertz (20). It also allows live video recording for later offline analysis. For this parabolic flight campaign, the effect of microgravity was studied on neuronal spheroids derived from a hippocampal neural progenitor grown under differentiating conditions in non-adherent cell culture plates. This neuronal progenitors spontaneously self-organize into spheres containing approximately 70% of hippocampal neurons among which 25% are GABAergic neurons, and 30% of astrocytes (Fig. 3). The typical diameter of a spheroid is about 200 *µm*. These 3D networks were selected because they allow the observation of spontaneous and synchronized calcium transients, providing information on the overall behavior of the spheroids without requiring external stimulation.

**Fig. 3.**
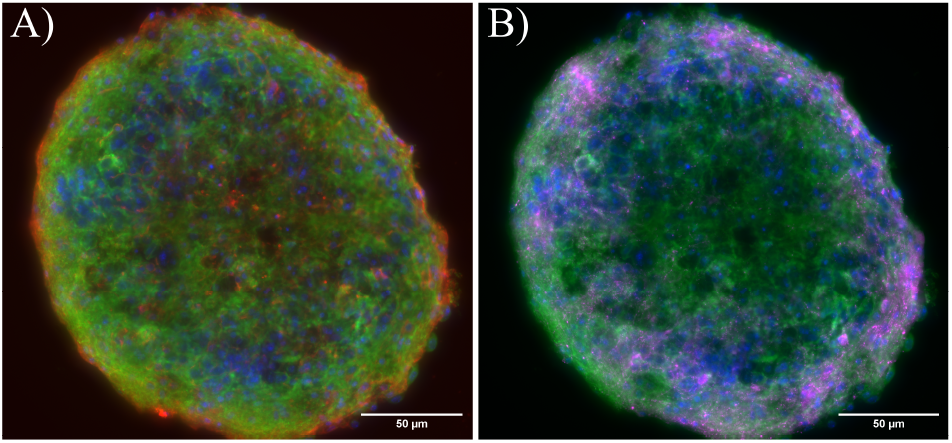
Production of neuronal spheroids. Immunostaining of a section of atypical 15 DIV hippocampal neuron spheroid grown for the flight campaign. A) staining of cell Nuclei (DAPI blue), Neuronal Dendrites (MAP2, green) and Astrocytes (GFAP, Red), B) staining of cell Nuclei (DAPI blue), Neuronal Dendrites (MAP2, green) and GABAergic neurons. (Gad67,magenta.) Both scale bars are 50 *µm*.

During two flights, the calcium activity of 20 active spheroids from 2 different cultures (i.e. different mice) was imaged and recorded. Fig. 4B shows a typical snapshot of a spheroid in acoustic levitation as monitored with the fluorescence microscope during the parabola. Each recording of a spheroid contains two parabolas, it starts 50 *s* before the onset of the first parabolas, i.e. before the hypergravity phase (1.8 *g*), and ends when the aircraft returns to 1 *g*, at the end of the second parabola. Each parabola consisted of approximately 30 *s* of hypergravity followed by 22 *s* of microgravity and finally ending with 30 *s* of hypergravity. Thus, each recording lasted approximately 300 *s*. The time evolution of the three components of the acceleration of the plane over two parabolas are plotted in the upper graph in Fig. 4A. The corresponding time evolution of calcic activity of a spheroid is shown in the two lower graphs in Fig. 4A. Throughout the campaign, approximately 15 *min* of calcium activity of primary hippocampal neurons was recorded in microgravity and more than 30 *min* in hypergravity.

**Fig. 4.**
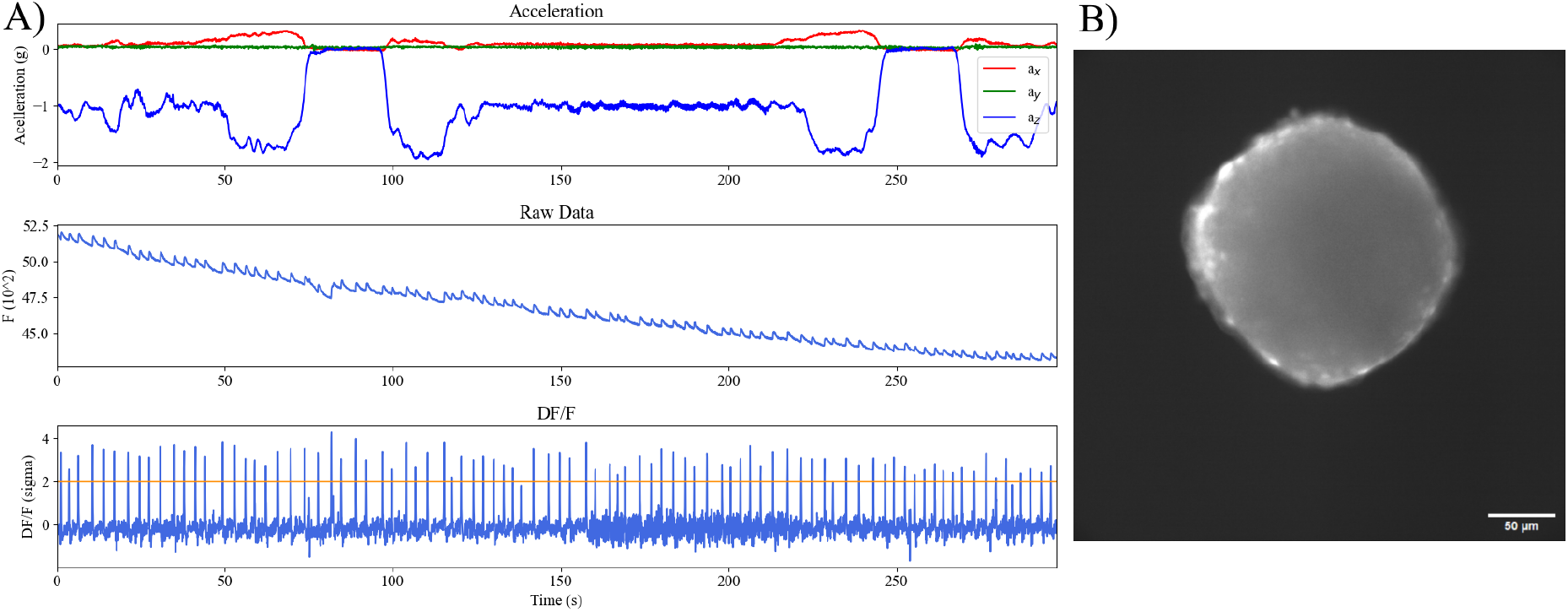
Activity of primary hippocampal neurons during parabolic flights. (A1) Acceleration on the three different axis in g (earth gravity 9.81*m*.*s*^*−*2^), during two parabolas plotted with respect to time. (A2) Typical fluorescent signal from calcic activity of neurons imaged during a parabola, 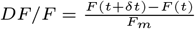where *F* (*t*) is the intensity of the spheroid for a given time and *F*_*m*_ the mean intensity value of the field of view. This value is then normalized by the standard deviation. The orange line represents the threshold value for a spike, which is defined as an amount that is two times the standard deviation. B) Typical view of a 15-days-old hippocampal spheroid stained with Fluo4 (ThermoFisher Scientific), scale bar is 50 *µm*.

### Influence of gravity changes on calcium activity of spheroids in acoustic levitation during parabolic flights

The activity of the spheroids was recorded during the three gravity phases (standard gravity, hypergravity and microgravity). The bursting period was extracted for all gravity phases during 20 parabolas. The mean values and the distribution of the activity periods are plotted as violins in Fig. 5. It varies from 5.49 *±* 3.51s in 1*g* to 4.75 *±* 3.34s in 1.8*g* and 3.16 *±* 2.56s in 0*g*. The first observation is that hypergravity does not induce a significant variation of the bursting period, only a small reduction of the mean period. On the contrary, microgravity causes a significant decrease of the mean bursting period. Moreover, the dispersion of the bursting periods is also reduced.

**Fig. 5.**
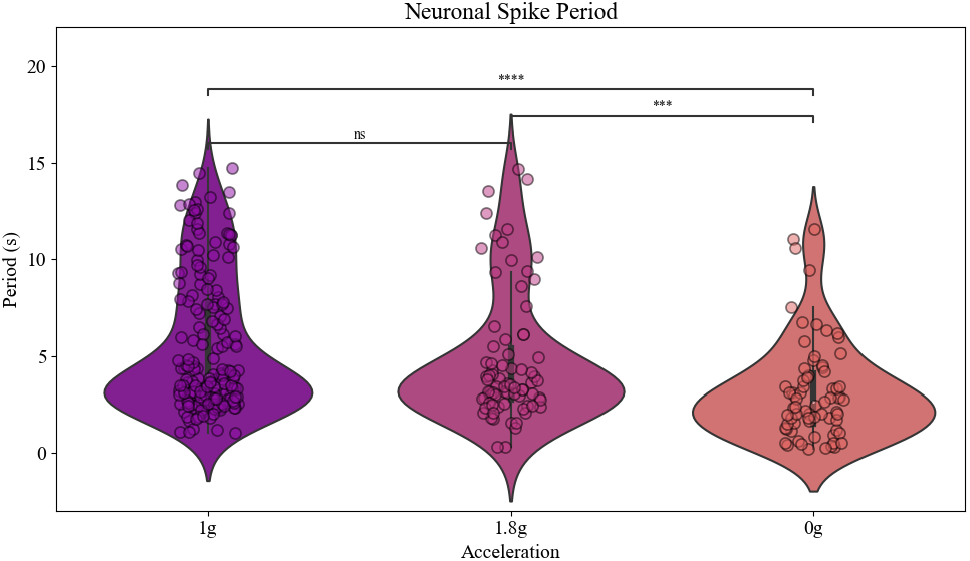
Calcium activity of neurospheres imaged during a parabola. Violin plots for comparison of firing period from hippocampal neurospheres, for each of the three phases of gravity. Mann-Whitney-Wilcoxon test two-sided were used (^*∗*^*p <* 0.05, ^*∗∗*^*p <* 0.01, ^*∗∗∗*^*p <* 0.001, and ^*∗∗∗∗*^*p <* 0.0001).

These first experiments demonstrate that acoustic levitation allows efficient trapping and rapid imaging of live neuronal spheroids during a 3-hour parabolic flight. Our results clearly show that mature neuronal networks respond specifically to microgravity by accelerating their internal rhythms.

Changes in neuronal activity due to microgravity has been postulated to be caused by modification of membrane viscosity and/or excitability (21). Changes in neuronal activity due to microgravity has been postulated to be caused by modification of membrane viscosity and/or excitability (21). We therefore wondered whether increasing neuronal spheroids excitability trough 4-Amino Pyridine treatment, a non-selective voltage-dependent 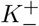channel blocker, would modify microgravity induced change in activity.

In a second set of experiments, 15 DIV hippocampal spheroids treated with 50*µ* M 4-aminopyridine (4-AP) were studied. As shown in Fig. 6, hippocampal spheroids exhibited faster bursting activity during the 1*g* phase compared to untreated spheroids. This is an expected result since it has been previously described that 4-AP significantly increases the bursting activity of 3D networks (22).

**Fig. 6.**
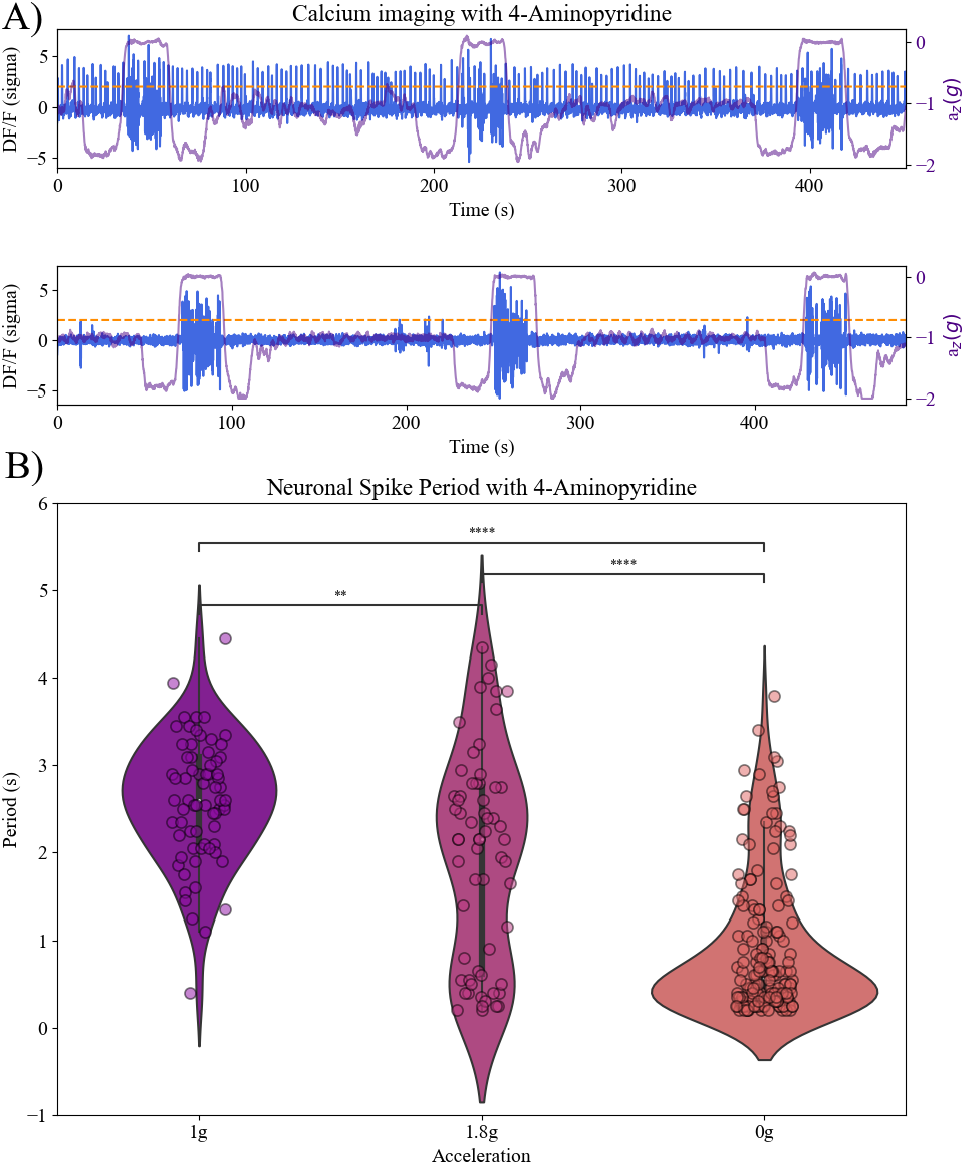
Calcium activity of neurospheres treated with 4-AP imaged during a parabola. (A) Two typical fluorescent signals from neurons treated with 4AP imaged during three consecutive parabolas, the first one is continuously active whereas the second is only active during the microgravity phase. 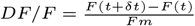where *F* (*t*) is the intensity of the spheroid for a given time and *Fm* the mean intensity value of the field of view, plotting as a function of standard deviation. In purple is the acceleration along the z axis plot as a function of the earth gravity (a_*z*_ (g)). (B) Violin plots of hippocampal neurospheres activity for each of the three phases of gravity. Mann-Whitney-Wilcoxon test two-sided were used (^*∗*^*p <* 0.05, ^*∗∗*^*p <* 0.01, ^*∗∗∗*^*p <* 0.001, and ^*∗∗∗∗*^*p <* 0.0001). The dotted line represents the spike detection threshold.

In comparison with the untreated spheroids (Fig. 5), the hypergravity phase modify the bursting period of the 4-APtreated spheroids compared to the 1*g* phase (Fig. 6). Strikingly, the microgravity phases led to a specific and dramatic decrease in the bursting period of the spheroids. We also observed a significant modification of the envelope of the violin distribution. Indeed the activity at 1*g* is slower than at 0*g* where it seems to reach a threshold at short period (which is for the fastest 4 times slower than the acquisition speed which seems to indicate that it is a plateau linked to the spheroid and not to the acquisition). The activity at 1.8*g* seems to be bimodal, with both average response times of 1*g* and 0*g*.

It has been shown that neural network oscillations occur through the fine-tuning of the excitatory/inhibitory balance, a process that occurs through a percolation scenario (23). The result is that at the time of sampling, some spheroids are spontaneously active while others are not. In our case, some of the 4-AP-treated spheroids showed bursting only at the beginning, in the 1*g* and 1.8*g* phases. Strikingly, in these cases, the spheroids woke up only during the microgravity phase. Two typical time-series of calcium activity variations during the three phases of gravity are shown in Fig. 6 where the first one represents an always active spheroid while the second only burst in microgravity.

Taken together our data strongly suggest that 3D neuronal spheroids specifically react to microgravity changes in a very specific manner presumably trough changes in intrinsic excitability of neurons.

## Discussion

With the advances in deep space exploration technologies and the increase in manned space missions, space biotechnologies play an increasingly important role, both to develop sustainable and efficient methods to increase the well-being of astronauts during space missions and to develop new technologies inaccessible on Earth (24, 25). In this context, the development of techniques for manipulating living cells, such as acoustic levitation, makes it possible to consider studies on the effects of spaceflight conditions on biological objects that are very difficult to study in microgravity. We have thus developed and validated an experimental device that allows the capture and trapping of 3D living objects, reminiscent of organoids, using acoustic tweezers, thus enabling live dynamic imaging of a rapid biological response. Here are some key interests and potential advantages of using acoustic levitation for space bioproduction.

As the manipulation of 3D cellular spheroids and organoids in weightlessness becomes a necessity (26), the present study demonstrates that acoustic waves allow a very efficient trapping of large living objects (hundreds of *µ*m) as they have already been used to trap nanorods in microgravity (12). Only a few experimental methods allow to manipulate 3D cellular objects in microgravity while allowing their observation. Besides hydrogel trapping and/or microfluidic constraint, magnetic levitation has been proposed as an interesting alternative since it allows to move cells in all 3 directions (11). However, magnetic levitation requires the magnetization of cells by labeling with nanoparticles and the use of paramagnetic cell culture media, a process that can be problematic for longterm studies (27, 28). In this context, acoustic levitation appears to be a promising alternative strategy, as it requires no cell labeling, no modification of the culture medium, and no mechanical contact.

Acoustic levitation is a promising tool for bioconstruction in space. It is based on the creation of a standing wave pattern leading to an acoustic radiation pressure field which, without contact, acts on all objects suspended in a fluid medium.

The acoustic forces act on particles by grouping and trapping them in disk-shaped layers (29, 30), located at the different pressure nodes of a resonant cylindrical cavity (31). It has already been used to form and culture cell spheroids of MSCs (Mesenchymatous Stromal Cells) or hepatocytes for several days (14, 15). Once acoustically levitated, objects (cell sheets, spheroids) can be further manipulated by variations in the magnitude of the force or by changing the distance between the acoustic pressure nodes (16, 17, 32, 33).

Acoustic levitation has been shown to maintain floating spheroids in a scaffold-free environment with precise control of axial position, enabling long-term, fast, inverted, fluorescent imaging of active neural spheroids. Significant, but limited, lateral movements of the spheroids were nevertheless observed during parabolic flights. These are caused by the acceleration of the *X* axis of the plane during the execution of the parabolas, and by the fact that the transverse ARF is 2 orders of magnitude smaller than the axial component of the ARF. However, this did not prevent calcium analysis of the spheroids as tracking algorithms can easily cope with that issue. It is important to note that drift in the levitation plane is limited to the specific conditions of parabolic flights during which hypergravity phases alternate with microgravity phases and is not supposed to occurs in constant 0g/microgravity environments such as the one observed in ISS. Importantly, the trapping efficiency can be easily controlled by changing the ARF amplitude. While this can mitigate the drift of objects in the levitation plane during parabolic flights, it could also be an interesting tool to modify the forces around levitating spheroids, simulate the variation of *g* in microgravity environments, and/or implement dynamic control of mechanical forces in cell culture experiments. In the present study, a simple, low-power, broadband piezoelectric transducer was used. Piezoelectric transducer arrays could be implemented in multiple directions, allowing for even more precise control of cells/spheroids in a 3D environment (34).

The use of these 3D cellular spheroids/organoids handling platforms makes it possible to study new, formerly inaccessible biological objects such as neuronal spheroids to mimic brain activity. Neuronal connectivity and brain activity are highly dynamic and interconnected processes, requiring detailed studies to understand how microgravity and space conditions impact the complex functions of neuron networks. This understanding is essential for addressing the effects of spaceflight on the nervous system. At the cellular level, microgravity has been consistently showed to directly affect cells, leading to fast modifications in their cytoskeleton and changes in the physical properties of their membranes but only a few experiments have been conducted until now (35– 38).

In this study, a decrease in the period of spontaneous activity of neuronal oscillations was observed during variations in gravity. The time between spikes is specifically reduced in microgravity compared to 1 *g* and 1.8 *g*. These results apparently contrast with a previous study which showed a tendency to decrease the firing of hippocampal neurons cultured in 2D networks during the 0*g* and 1.8*g* phases(39). Yet the experiment was designed to study the impact of altered gravity on single, unconnected, immature neurons. It showed that only a subset of cells showed a reduction in their firing rate, while a separate silent population began to burst, providing evidence that the effect of microgravity on the neuronal subpopulation could be twofold.. Here, we leveraged the use of neuronal spheroids that have specific advantages over immature brain organoids. Indeed, neuronal spheroids were fully mature and exhibited highly coordinated network activity, a result of their intrinsic excitatory/inhibitory balance (40).

Furthermore, the presented results are consistent with other experiments using measurements of activities of mature human iPSC-derived neurons with Micro-Electrode Array (MEA) recording platforms that indicated a slight increase in the frequency of network bursting during the 0*g* and 1.8*g* phases in a drop tower configuration (41). These results suggest that microgravity can have subtle effects on a single cell that result in entire networks adapting toward hyperactivity. The presented results show that promoting network hyperexcitability with a *K*^+^ channel blocker results in a dramatic increase in rapid bursting events, particularly during 0g phases, suggesting that microgravity has subtle but significant effects on cell membrane polarization. Although the molecular players mediating this effect remain elusive, a plausible explanation could be that microgravity induced changes in membrane tension and/or fluidity, leading to limited activation of ion channels in neuronal membranes, a process revealed by pharmacological hyperexcitability. Indeed, membrane deformation can modify the function of mechanoreceptors, some of which participate in membrane polarization (42).

Indeed, it is known that a single mechanical indentation localized in the *pN* range can cause a local and transient calcium response in (43) cells. It is therefore tempting to speculate that microgravity triggers sub threshold activation of ion mechanoreceptors, which, in conjunction with 4-AP, leads to rapid depolarization. An alternative but not exclusive hypothesis being that the microgravity phase modifies the probability of presynaptic glutamate release or the content of postsynaptic glutamatergic receptors. Indeed, although this has not been formally tested in a real microgravity environment, artificial gravity experiments have shown changes in glutamatergic pathways (44–46). Finally, the spheroids having been observed under acoustic trapping, the ultrasound itself could participate in the observed effects. Indeed, low intensity ultrasound is known to modulate neuronal excitability through the activation of ionic mechanoreceptors (47, 48) and common molecular signals could theoretically be involved in an alteration of gravity and/or effects biological ultrasound (49).

The intrinsically constrained experimental environment of parabolic flight represents an important challenge for studying the precise molecular and cellular mechanisms underlying the subtle effect of microgravity on the excitability of neuronal membranes and how networks self -adapt to these modifications. Further research at higher throughput is needed to address this challenge.

With a growing need to study the consequences of the space environment on human physiology and to design new means to ensure survival in space, including bioproduction in space, manipulation of complex cellular systems in a microgravity environment becomes an important issue. Therefore, experimental tools to manipulate and interrogate 3D structures such as organoids are needed (50). Thanks to their excellent biocompatibility and versatility, acoustic approaches are promising tools for contactless dynamic manipulation of living cells on Earth and in space (14, 15, 51). Thanks to the fine tuning of parameters at the origin of the ARF, such as the acoustic frequency, or the use of networks of transducers to control the spatial organization of the acoustic field, the manufacturing and manipulation of complex 3D cellular assemblies are possible (52, 53). By levitating these biological samples in a liquid medium, it is possible to observe their behaviors and interactions without the influence of gravity, thus providing valuable information for space biology experiments carried out aboard platforms such as the International Space Station (ISS) (11). This technique can address some of the unique challenges associated with cell culture, bioproduction or tissue engineering in microgravity and harsh extraterrestrial environments.

## Materials and Methods

### A. Acoustofluidic setup

The micro chip were composed of a PDMS (Polydimethylsiloxane) layer bonded on top of a microscope cover-glass and further connected to a mono-element ultrasonic transducer. The design of the microfluidic chips was coded on a brass mold generated with a micromilling machine. To fabricate the chips, a mixture of PDMS elastomer and curing agent in a ratio of 9:1 was degassed under vacuum to remove air bubbles. The PDMS mixture was then poured onto a mold and baked for 4 hours at 70°C. The PDMS microfluidic chips were peeled off from the mold, and the PDMS surface was cleaned with adhesive tape. The PDMS buffers were then stuck on glass slides, previously cleaned with isopropanol as well as using a plasma cleaner (power 100%, 0.6 *mBar* in *O*_2_, Diener Electronic). The chamber was immediately filled with distilled water maintaining the hydrophilicity of glass and PDMS acquired during plasma treatment. Finally, the microfluidic chips were sterilized under UV for 30 min.

The geometry of the PDMS chip is designed to allow insertion of the ultrasonic transducer on top of the PDMS body. The transducer closes the acoustofluidic cavity. One side of the resonant cavity is the ultrasound emitter, while on the other side the microscope cover plate acts as an effective acoustic reflector and allows optical access. The overall acoustic resonant cavity is simple and effective. Ultrasound waves were generated by a transducer driven by an arbitrary waveform generator (Handyscope HS5 from Tiepie Engineering, Sneek, The Netherlands) monitored by a computer. Each output of the signal generators fed two ultrasonic transducers (2 *MHz* SignalProcessing, Savigny, Switzerland) with a sinusoidal waveform of amplitude 5*V*_*pp*_ and frequency 2 *MHz*. The acoustic wave parameters were chosen to optimize the levitation process while avoiding unwanted phenomena such as acoustic streaming.

### B. Cell culture of Hippocampal spheroids

Hippocampal neuronal cultures were generated from E18 SWISS mouse embryos by building upon standard cell-culture techniques, as follows. SWISS pregnant mice were purchased from Janvier (Le Genest Saint Isle, France). Animal care was conducted in accordance with standard ethical guidelines. The cell culture preparation was done in the Neurocentre Magendie laboratory in Bordeaux. Hippocampi were micro-dissected from E18 embryos with all steps performed in cold Gey’s Balanced Saline Solution (Sigma G9779). Dissected structures were digested with papaïn (50 U/mL, Sigma 76220) in DMEM (Thermofisher 31966 021). After papaïn inactivation with FBS (100 U/mL, GE Healthcare), structures were mechanically dissociated with a pipette (plastic tip 1 mm diameter) in presence of DNAse type IV (20U/mL, Sigma D5025). Cells viability was determined by Trypan Blue exclusion assay. The resulting cells were centrifuged 7 *min* at 800 *rpm* and the pellet was resuspended in the culture medium containing DMEM, 10% FBS, 2% B27, 0.5mM Glutamate and 1% Penicillin/Streptomycin (Gibco^®^) at a concentration of 20 million cells/mL. Cells were seeded in microwells of U-shaped low adhesion plates (Nunclon Sphera, Thermofischer). All cell cultures were kept in an incubator at 37°C and 5% CO2. Spheroids were cultured 14 days before transfering to the Novespace biological facility laboratory for the 3 days of the flight campaign. The typical diameter of the spheroids obtained was around 200 *µm*.

### C. Calcium Imaging

The calcium dye Fluo4-AM was obtained from ThermoFisher Scientific^®^ (ref F14201). It was dissolved in dimethyl sulfoxide (DMSO) in a ratio of 1 *µg/µL* to obtain a stock solution of 1 mM. This stock solution was frozen in 2 *µ*L aliquots. *Ca*^2+^ free wash buffer was made according to previous protocols and the stock solution of Fluo4-AM was added in order to load the cells with a final concentration of approximately 2 *µ*M. Before the flights or classical on-ground experiments, spheroids were chosen, loaded in a microfluidic chip with Fluo 4 AM and incubated at 37^*o*^C and 5% CO_2_ for 25 min. Spheroids were then placed in culture medium (brainphy) with 50*µM* of bicuculine (Sigma-Aldrich ref 505875). For the 4-AP treatments, spheroids were similarly treated with 50*µM* of 4-AP. Finally, after sealing the inlet and outlet reservoirs with a PDMS membrane to mitigate fluid movement during flight, the chips were transported in a closed box (to protect the cells from light as well as temperature variations) On board the plane. The equipment is described in the results section below, the exposure time was 20 *ms* every 50 *ms*. The activity recorded was purely spontaneous for all measurements. For analysis, the signal is normalized using 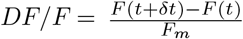 where *F* (*t*) is the spheroid intensity for a given time and *F*_*m*_ the average value of the intensity of the field of vision. From this normalized signal, only values exceeding 2 times the standard deviation are counted as bursts. Average firing and burst rates were extracted. Bursting is a phenomenon observed in neurons and is characterized by a brief increase in signal followed by a longer return to the initial point. The Mann–Whitney U test was used for statistical analysis. All analyzes and plots were generated using Python routines.

### D. Experimental setup adapted to Zero-G flights

ssTo perform the calcium imaging experiments in the Airbus ZeroG aircraft, a previously modified setup dedicated to calcium imaging of living cells was used (39). In short, the setup is made up of three modules. The main new constraints of the device were linked to the manipulation and observation of living cells by fluorescence microscopy in a very confined environment. A monitoring rack was designed to allow control of key experimental parameters and rapid data acquisition with two computers, each recording images from different experiments. The experimental setup (Fig. 7) was designed to fit in a sealed Zarges box for safety reasons. Acoustically levitating particles and cells were studied using a fully automated Zeiss Axiobserver inverted microscope equipped with a stage incubator (Tokai Hit), an LED array (Coolled pE 4000) and equipped with a Hamamastu Orca Fusion CMOS camera. Aggregates of particles or cells can be observed with small DinoLite cameras placed on a 3D printed mount. This allows the behavior of acoustically levitating spheroids or beads to be observed from the side.

**Fig. 7.**
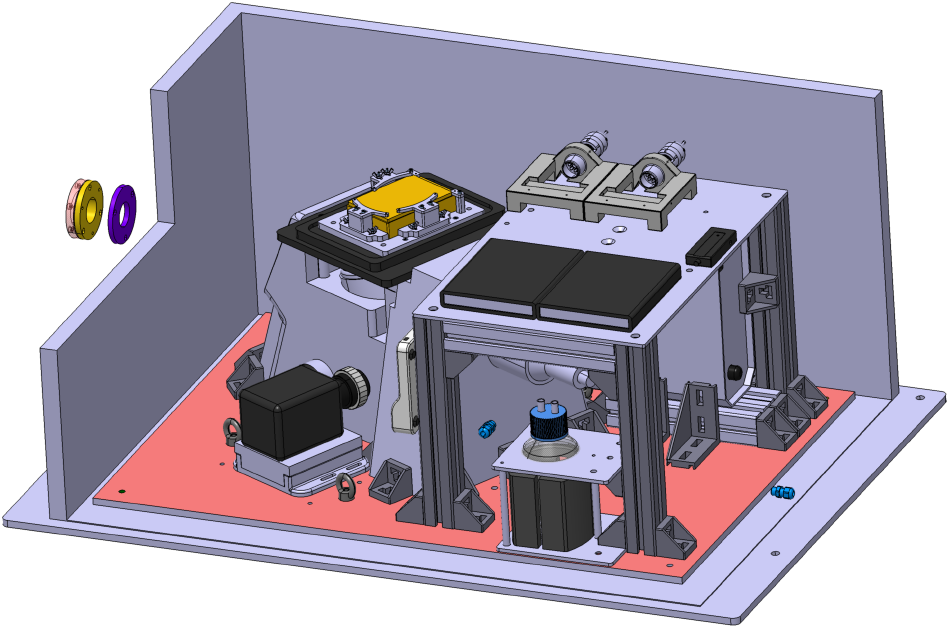
3D view of the experimental box. (Rack 2). Inside a sealed Zarges box, we installed a Zeiss inverted microscope with a motorized stage. A stage incubator from TOKAI HIT was used to ensure a well-controlled environment for the cell cultures during the flights. The fluorescence images were recorded using a Hamamastu Orca Fusion camera.

## ACKNOWLEDGEMENTS

We thank the CNES (Centre National d’Etudes Spatiales) for its financial support (APR CNES). We also thank C.Delaroche and T. Bret-Dibat (topic “Sciences de la Matière”) and G. Gauquelin-Koch (topic “Sciences de la Vie dans l’Espace”) who made possible our participation to the VP177 Parabolic Flights campaign. We also thank the Novespace team, and especially Thibault Paris, for helping us to properly take into account the many constraints imposed by zero-g flights, making these experiments possible. We warmly thank Aude Panatier and the Neurocentre Magendie (INSERM, University of Bordeaux) who allowed us to access their laboratory in order to prepare and cultivate the cells on site. We would also like to thank the Institut Pierre-Gilles de Gennes and in particular Audric Jan for micromilling the brass microfluidic molds.

## Bibliography

1. K. E. Hupfeld, H. R. McGregor, P. A. Reuter-Lorenz, and R. D. Seidler. Microgravity effects on the human brain and behavior: Dysfunction and adaptive plasticity. Neuroscience & Biobehavioral Reviews, 122:176–189, March 2021. ISSN 0149-7634. doi: 10.1016/j.neubiorev.2020.11.017.

2. Udit Gupta, Sheharyar Baig, Arshad Majid, and Simon M Bell. The neurology of space flight; How does space flight effect the human nervous system? Life Sciences in Space Research, September 2022. ISSN 2214-5524. doi: 10.1016/j.lssr.2022.09.003.

3. Meaghan Roy-O’Reilly, Ajitkumar Mulavara, and Thomas Williams. A review of alterations to the brain during spaceflight and the potential relevance to crew in long-duration space exploration. npj Microgravity, 7(1):1–9, February 2021. ISSN 2373-8065. doi: 10.1038/s41526-021-00133-z. Number: 1 Publisher: Nature Publishing Group.

4. Jessica K. Lee, Vincent Koppelmans, Roy F. Riascos, Khader M. Hasan, Ofer Pasternak, Ajitkumar P. Mulavara, Jacob J. Bloomberg, and Rachael D. Seidler. Spaceflight-Associated Brain White Matter Microstructural Changes and Intracranial Fluid Redistribution. JAMA Neurology, 76(4):412–419, April 2019. ISSN 2168-6149. doi: 10.1001/jamaneurol.2018.4882.

5. Steven Jillings, Ekaterina Pechenkova, Elena Tomilovskaya, Ilya Rukavishnikov, Ben Jeurissen, Angelique Van Ombergen, Inna Nosikova, Alena Rumshiskaya, Liudmila Litvinova, Jitka Annen, et al. Prolonged microgravity induces reversible and persistent changes on human cerebral connectivity. Communications Biology, 6(1):46, 2023.

6. Francesca Ferranti, Marta Del Bianco, and Claudia Pacelli. Advantages and limitations of current microgravity platforms for space biology research. Applied Sciences, 11(1):68, 2020.

7. Alexa Wnorowski, Arun Sharma, Haodong Chen, Haodi Wu, Ning-Yi Shao, Nazish Sayed, Chun Liu, Stefanie Countryman, Louis S. Stodieck, Kathleen H. Rubins, Sean M. Wu, Peter H.U. Lee, and Joseph C. Wu. Effects of spaceflight on human induced pluripotent stem cell-derived cardiomyocyte structure and function. Stem Cell Reports, 13(6):960–969, 2019. ISSN 2213-6711. doi: 10.1016/j.stemcr.2019.10.006.

8. Carlos Cepeda, Laurent Vergnes, Nicholas Carpo, Matthew J Schibler, Laurent A Bentolila, Fathi Karouia, and Araceli Espinosa-Jeffrey. Human neural stem cells flown into space proliferate and generate young neurons. Applied Sciences, 9(19):4042, 2019.

9. Lorenzo Moroni, Kevin Tabury, Hilde Stenuit, Daniela Grimm, Sarah Baatout, and Vladimir Mironov. What can biofabrication do for space and what can space do for biofabrication? Trends in Biotechnology, 40(4):398–411, 2022.

10. Ruimin Long, Linrong Shi, Peng He, Jumei Tian, Shibin Wang, and Jun Zheng. 3d cell culture based on artificial cells and hydrogel under microgravity for bottom-up microtissue constructs. Frontiers in Bioengineering and Biotechnology, 10:1056652, 2022.

11. Vladislav A Parfenov, Yusef D Khesuani, Stanislav V Petrov, Pavel A Karalkin, Elizaveta V Koudan, Elizaveta K Nezhurina, Frederico DAS Pereira, Alisa A Krokhmal, Anna A Gryadunova, Elena A Bulanova, et al. Magnetic levitational bioassembly of 3d tissue construct in space. Science advances, 6(29):eaba4174, 2020.

12. Gabriel Dumy, Nathan Jeger-Madiot, Xavier Benoit-Gonin, Thomas E Mallouk, Mauricio Hoyos, and Jean-Luc Aider. Acoustic manipulation of dense nanorods in microgravity. Microgravity Science and Technology, 32:1159–1174, 2020.

13. Glenn Whitworth and WT Coakley. Particle column formation in a stationary ultrasonic field. The Journal of the Acoustical Society of America, 91(1):79–85, 1992.

14. Nathan Jeger-Madiot, Lousineh Arakelian, Niclas Setterblad, Patrick Bruneval, Mauricio Hoyos, Jérôme Larghero, and Jean-Luc Aider. Self-organization and culture of mesenchymal stem cell spheroids in acoustic levitation. Scientific reports, 11(1):8355, 2021.

15. Lucile Rabiet, Lousineh Arakelian, Nathan Jeger-Madiot, Duván Rojas García, Jérôme Larghero, and Jean-Luc Aider. Acoustic levitation as a tool for cell-driven self-organization of human cell spheroids during long-term 3d culture. Biotechnology and Bioengineering, 121(4):1421–1433, 2024.

16. Olivier Dron and Jean-Luc Aider. Varying the agglomeration position of particles in a micro-channel using acoustic radiation force beyond the resonance condition. Ultrasonics, 53(7):1280–1287, 2013.

17. Nathan Jeger-Madiot, Xavier Mousset, Chloé Dupuis Lucile Rabiet, Mauricio Hoyos, Jean-Michel Peyrin, and Jean-Luc Aider. Controlling the force and the position of acoustic traps with a tunable acoustofluidic chip: Application to spheroid manipulations. The Journal of the Acoustical Society of America, 151(6):4165–4179, 2022.

18. Stefania Torino, Brunella Corrado, Mario Iodice, and Giuseppe Coppola. Pdms-based microfluidic devices for cell culture. Inventions, 3(3):65, 2018.

19. G. Bradski. The OpenCV Library. Dr. Dobb’s Journal of Software Tools, 2000.

20. Jean-Pierre Eckmann, Ofer Feinerman, Leor Gruendlinger, Elisha Moses, Jordi Soriano, and Tsvi Tlusty. The physics of living neural networks. Physics Reports, 449(1):54–76, 2007. ISSN 0370-1573. doi: 10.1016/j.physrep.2007.02.014. Nonequilibrium physics: From complex fluids to biological systems III. Living systems.

21. FPM Kohn and R Ritzmann. Gravity and neuronal adaptation, in vitro and in vivo-from neuronal cells up to neuromuscular responses: a first model. Eur Biophys J., 2018. doi: 10.1007/s00249-017-1233-7.

22. Alfredo Gonzalez-Sulser, Jing Wang, Bridget N Queenan, Massimo Avoli, Stefano Vicini, and Rhonda Dzakpasu. Hippocampal neuron firing and local field potentials in the in vitro 4-aminopyridine epilepsy model. Journal of neurophysiology, 108(9):2568–2580, 2012.

23. Ilan Breskin, Jordi Soriano, Elisha Moses, and Tsvi Tlusty. Percolation in living neural networks. Physical review letters, 97(18):188102, 2006.

24. Nils JH Averesch, Aaron J Berliner, Shannon N Nangle, Spencer Zezulka, Gretchen L Vengerova, Davian Ho, Cameran A Casale, Benjamin AE Lehner, Jessica E Snyder, Kevin B Clark, et al. Microbial biomanufacturing for space-exploration—what to take and when to make. Nature Communications, 14(1):2311, 2023.

25. Aaron J Berliner, Isaac Lipsky, Davian Ho, Jacob M Hilzinger, Gretchen Vengerova, Georgios Makrygiorgos, Matthew J McNulty, Kevin Yates, Nils JH Averesch, Charles S Cockell, et al. Space bioprocess engineering on the horizon. Communications Engineering, 1(1):13, 2022.

26. Davide Marotta, Laraib Ijaz, Lilianne Barbar, Madhura Nijsure, Jason Stein, Twyman Clements, Jana Stoudemire, Paula Grisanti, Scott A Noggle, Jeanne F Loring, et al. Studies on the international space station to assess the effects of microgravity on ipsc-derived neural organoids. bioRxiv, pages 2023–08, 2023.

27. Naside Gozde Durmus, H Cumhur Tekin, Sinan Guven, Kaushik Sridhar, Ahu Arslan Yildiz, Gizem Calibasi, Ionita Ghiran, Ronald W Davis, Lars M Steinmetz, and Utkan Demirci. Magnetic levitation of single cells. Proceedings of the National Academy of Sciences, 112 (28):E3661–E3668, 2015.

28. Alexandre Fromain, Aurore Van de Walle, Guilhem Curé, Christine Péchoux, Aida Serrano, Yoann Lalatonne, Ana Espinosa, and Claire Wilhelm. Biomineralization of magnetic nanoparticles in stem cells. Nanoscale, 15(23):10097–10109, 2023.

29. Jose P Leao-Neto, Mauricio Hoyos, Jean-Luc Aider, and Glauber T Silva. Acoustic radiation force and torque on spheroidal particles in an ideal cylindrical chamber. The Journal of the Acoustical Society of America, 149(1):285–295, 2021.

30. Mikkel Settnes and Henrik Bruus. Forces acting on a small particle in an acoustical field in a viscous fluid. Physical Review E, 85(1):016327, 2012.

31. Henrik Bruus. Theoretical microfluidics, volume 18. Oxford university press, 2007.

32. Martin Wiklund. Acoustofluidics 12: Biocompatibility and cell viability in microfluidic acoustic resonators. Lab on a Chip, 12(11):2018–2028, 2012.

33. Reza Rasouli, Radu Alexandru Paun, and Maryam Tabrizian. Sonoprinting nanoparticles on cellular spheroids via surface acoustic waves for enhanced nanotherapeutics delivery. Lab on a Chip, 23(8):2091–2105, 2023.

34. Bruce W Drinkwater. A perspective on acoustical tweezers—devices, forces, and biomedical applications. Applied Physics Letters, 117(18), 2020.

35. Susan J Crawford-Young. Effects of microgravity on cell cytoskeleton and embryogenesis. International journal of developmental biology, 50, 2006.

36. M Janmaleki, M Pachenari, SM Seyedpour, R Shahghadami, and A Sanati-Nezhad. Impact of simulated microgravity on cytoskeleton and viscoelastic properties of endothelial cell. Scientific reports, 6(1):32418, 2016.

37. Florian PM Kohn and Jens Hauslage. The gravity dependence of pharmacodynamics: the integration of lidocaine into membranes in microgravity. npj Microgravity, 5(1):5, 2019.

38. Yi Cui, Jin Han, Zhifeng Xiao, Yiduo Qi, Yannan Zhao, Bing Chen, Yongxiang Fang, Sumei Liu, Xianming Wu, and Jianwu Dai. Systematic analysis of mrna and mirna expression of 3d-cultured neural stem cells (nscs) in spaceflight. Frontiers in Cellular Neuroscience, 11:434, 2018.

39. Pierre-Ewen Lecoq, Chloé Dupuis Mousset Xavier, Xavier Benoit-Gonin, Jean-Michel Peyrin, and Jean-Luc Aider. Influence of microgravity on spontaneous calcium activity of primary hippocampal neurons grown in microfluidic chips. npj Microgravity, 2024. doi: 10.1038/s41526-024-00355-x.

40. Jessica L Sevetson, Brian Theyel, and Diane Hoffman-Kim. Cortical spheroids display oscillatory network dynamics. Lab on a Chip, 21(23):4586–4595, 2021.

41. Johannes Striebel, Laura Kalinski, Maximilian Sturm, Nils Drouvé, Stefan Peters, Yannick Lichterfeld, Rouhollah Habibey, Jens Hauslage, Sherif El Sheikh, Volker Busskamp, et al. Human neural network activity reacts to gravity changes in vitro. Frontiers in Neuroscience, 17:1085282, 2023.

42. Weikang Wang, Kuan Tao, Jing Wang, Gen Yang, Qi Ouyang, Yugang Wang, Lei Zhang, and Feng Liu. Exploring the inhibitory effect of membrane tension on cell polarization. PLoS computational biology, 13(1):e1005354, 2017.

43. Fabio Falleroni, Ulisse Bocchero, Simone Mortal, Yunzhen Li, Zhongjie Ye, Dan Cojoc, and Vincent Torre. Mechanotransduction in hippocampal neurons operates under localized low piconewton forces. Iscience, 25(2), 2022.

44. Gyutae Kim and Kyu-Sung Kim. Hypergravity-induced malfunction was moderated by the regulation of nmda receptors in the vestibular nucleus. Scientific Reports, 11(1):17420, 2021.

45. T Borisova, N Krisanova, and N Himmelreich. Exposure of animals to artificial gravity conditions leads to the alteration of the glutamate release from rat cerebral hemispheres nerve terminals. Advances in Space Research, 33(8):1362–1367, 2004.

46. Yun Wang, Javed Iqbal, Yahui Liu, Rui Su, Song Lu, Guang Peng, Yongqian Zhang, Hong Qing, and Yulin Deng. Effects of simulated microgravity on the expression of presynaptic proteins distorting the gaba/glutamate equilibrium–a proteomics approach. Proteomics, 15 (22):3883–3891, 2015.

47. Benjamin Clennell, Tom GJ Steward, Kaliya Hanman, Tom Needham, Janette Benachour, Mark Jepson, Meg Elley, Nathan Halford, Kate Heesom, Eunju Shin, et al. Ultrasound modulates neuronal potassium currents via ionotropic glutamate receptors. Brain Stimulation, 16(2):540–552, 2023.

48. Sangjin Yoo, David R Mittelstein, Robert C Hurt, Jerome Lacroix, and Mikhail G Shapiro. Focused ultrasound excites cortical neurons via mechanosensitive calcium accumulation and ion channel amplification. Nature communications, 13(1):493, 2022.

49. Emilie Campanac, Célia Gasselin, Agnès Baude, Sylvain Rama, Norbert Ankri, and Dominique Debanne. Enhanced intrinsic excitability in basket cells maintains excitatory-inhibitory balance in hippocampal circuits. Neuron, 77(4):712–722, 2013.

50. Nicolette A Pirjanian, Kriti Kalpana, Ilya Kruglikov, Pinar Mesci, Jana Stoudemire, Paula Grisanti, Scott A Noggle, Jeanne F Loring, and Valentina Fossati. Establishing neural organoid cultures for investigating the effects of microgravity in low-earth orbit (leo). Springer, 2024.

51. K Olofsson, B Hammarström, and M Wiklund. Ultrasonic based tissue modelling and engineering. micromachines 9, 594, 2018.

52. Alexa Wnorowski, Huaxiao Yang, and Joseph C Wu. Progress, obstacles, and limitations in the use of stem cells in organ-on-a-chip models. Advanced drug delivery reviews, 140: 3–11, 2019.

53. Thomas M Llewellyn-Jones, Bruce W Drinkwater, and Richard S Trask. 3d printed components with ultrasonically arranged microscale structure. Smart Materials and Structures, 25 (2):02LT01, 2016.

